# Dysfunctional Senescent Herpes Simplex Virus Specific CD57^+^CD8^+^ T cells are Associated with Recurrent Symptomatic Herpes in Humans

**DOI:** 10.1101/2022.01.05.475169

**Authors:** Arif A. Khan, Ruchi Srivastava, Hawa Vahed, Lbachir BenMohamed

## Abstract

Herpes simplex virus (HSV)-specific CD8^+^ T cells protect mice from herpes infection and disease. However, the phenotype and function of HSV-specific CD8^+^ T cells that play a key role in the “natural” protection seen in HSV-1-seropositive healthy asymptomatic (ASYMP) individuals (who have never had clinical herpes disease) remain to be determined. We previously reported that symptomatic (SYMP) patients (who have frequent bouts of recurrent herpes disease) had more less-differentiated and dysfunctional HSV-specific CD8^+^ T cells. In contrast, healthy ASYMP individuals maintained a significantly higher proportion of differentiated polyfunctional CD8^+^ T cells. Here we report that, HSV-specific CD8^+^ T cells from SYMP patients, but not from ASYMP individuals, have phenotypic and functional characteristics of cellular senescence, including: (*i*) high frequency of senescent (CD57^+^) and exhausted (PD-1^+^) CD8^+^ T cells; (*ii*) late terminally differentiated (KLRG1^+^), non-proliferating CD8^+^ T cells; (*iii*) HSV-specific CD8^+^ T cells were declined overtime and were not maintained homeostatistically (CD127^+^CD8^+^ T cells); (*iv*) loss of co-stimulatory molecule (CD28) on HSV-specific CD8^+^ T cells; (*v*) decreased production of effector molecules (granzyme B and perforin) by HSV-specific CD8^+^ T cells. Our findings provide insights into the role of senescence in HSV-specific CD8^+^ T cells in susceptibility to recurrent herpes and have implications for T-cell-based immunotherapeutic strategies against recurrent herpes in humans.

**IMPORTANCE:** Reactivation of herpes simplex virus 1 (HSV-1) from latency leads to recurrent herpetic disease in symptomatic (SYMP) individuals. A role of T cells in herpes immunity against recurrent herpetic disease is established. However, the phenotype and function of HSV-specific CD8^+^ T cells that play a key role in the “natural” protection seen in HSV-1-seropositive healthy asymptomatic (ASYMP) individuals (who have never had clinical herpes disease) remain to be determined. In the present study, we report that, HSV-specific CD8^+^ T cells from SYMP patients, but not from ASYMP individuals, have phenotypic and functional characteristics of cellular senescence. Our findings provide insights into the role of senescence in HSV-specific CD8^+^ T cells in susceptibility to recurrent herpes and have implications for T-cell-based immunotherapeutic strategies against recurrent herpes in humans.

## INTRODUCTION

Over a billion individuals worldwide carry Herpes Simplex Virus type 1 (HSV-1) which causes a wide range of mild to life-threatening diseases (1, 2). Although the virus reactivates from latency and is shed multiple times each year in body fluids (i.e., tears, saliva, nasal, and vaginal secretions), most reactivations are subclinical due to an efficient immune-mediated containment of the infection and disease (3–6). Thus, most infected individuals are asymptomatic (ASYMP) and do not present any apparent recurrent herpetic disease (e.g., cold sores, genital, or ocular herpetic disease). However, a small proportion of individuals experience endless recurrences of herpetic disease, usually multiple times a year, often necessitating continuous antiviral therapy (i.e., acyclovir and derivatives) (7, 8). In those symptomatic (SYMP) individuals, HSV-1 frequently reactivates from latency, re-infects the eyes, and may trigger recurrent and severe corneal herpetic disease, a leading cause of infectious corneal blindness in the industrialized world (9–11). In the United States, up to 450,000 individuals have a history of recurrent herpetic stromal keratitis (rHSK), a T cell mediated immunopathological lesion of the cornea (9–11). Thus, a better understanding of the immune mechanisms that protect from HSV-1 infection and disease is highly desirable for the development of more efficacious vaccines and immunotherapies to reduce HSV-1 related diseases.

In ASYMP individuals, what are the factors that influence reactivation of the virus without any disease throughout lifespan remains an active field of investigation. The question, which we wanted to address in this report, is there any difference in the maintenance of optimal CD8^+^ T cell response between HSV-1 seropositive ASYMP and SYMP individuals. There are reports indicating that immunosenescence is marked by a progressive increase in the number of memory CD8^+^ T cells showing poor functionality in terms of killing persistent virus (12). We hypothesize that the long-term, repeated stimulation of the CD8^+^ T cells by reactivating herpes virus might lead to immunosenescence in SYMP individuals but not in ASYMP individuals. We explored the association of specific CD8^+^ T cell markers of immune differentiation (CD28), senescence (KLRG-1 and CD57) and survival of HSV-specific CD8^+^ T cells. In literature, CD57^+^CD8^+^ T cells have been shown to be associated with various inflammatory diseases, HIV, and chronic infection (13–17). However pathogenic roles of CD57^+^CD8^+^ T cells have not yet been elucidated in HSV-1 seropositive SYMP individuals. This study is a step forward where we assessed the immunological characteristics of CD57^+^CD8^+^ T cells in ASYMP and SYMP individuals.

The differentiation of T cell towards a senescent phenotype is characterized typically by clonal expansion of CD8^+^ T cells that lack expression of CD28 (13, 18). Recently it has been shown that CD8^+^CD28^-^CD57^+^ T cells fill the vital immunological space and render it inaccessible to the naive and early memory T-cell repertoire in early-stage HIV infection with more rapid progression to AIDS (13, 14, 16, 19). Expression of KLRG1 identifies CD8^+^ T cells in humans that are capable of secreting cytokines but fails to proliferate after stimulation and is thus unable to undergo further clonal expansion (20, 21). Increased frequency of late-senescent CD8^+^ T cells lacking survival marker CD127 (IL-7R) results in defective killing abilities and less surface expression of CD28 leads to defective proliferation abilities. Here, we demonstrate that increased percentage of CD8^+^ T cells from SYMP individuals express CD57 and KLRG1 and decreased percentage of CD28 and CD127. Loss of CD127 in HSV-specific CD8^+^ T cells could lead to apoptosis and thus lack homeostatic maintenance. The T-box transcription factor T-bet plays crucial roles in determining differential fate of CD8^+^ T cells responding to infection, optimal memory, and terminal differentiation. In CD8^+^ T cells, T-bet is up regulated upon activation and is associated with induction of effector functions, including cytotoxicity. Expression of T-bet was increased in CD8^+^ T cells from SYMP individuals and correlated closely with expression of CD57 and KLRG1. Although senescent CD8^+^ T cells typically have preserved effector function (i.e., the ability to produce cytokines and even kill target cells), lack of proliferative potential impairs their ability to mount robust immune responses and expand in number upon reactivation. During HIV infection, senescent CD57^+^CD8^+^ T cells produce IFN-γ but proliferate poorly and are more sensitive to activation-induced cell death. Thus, the accumulation of senescent CD8^+^ T cells could lead to deterioration of protective immune responses.

In this report we found that in HSV-1 seropositive SYMP individuals there was increased frequency of senescent CD8^+^ T cells, which lack co-stimulatory molecule CD28 and survival molecule CD127 as compared to ASYMP individuals. We also showed that higher expression of transcription factor T-bet was associated with increased co-expression of CD57 and KLRG1. Thus, cellular senescence appears to be a major feature of HSV-specific CD8^+^ T cells in the SYMP individuals and T-bet is likely involved in the underlying molecular regulation of this terminally differentiated state. Altogether, these results indicate that, there is significantly higher frequency of HSV specific senescent CD8^+^ T cell in SYMP individuals, which lack proliferative capacity, co-stimulatory marker and cannot maintain homeostatic proliferation as compared to HSV-seropositive ASYMP individuals. These findings have implications for T-cell-based immunotherapeutic strategies designed for recurrent herpes in humans

## MATERIALS AND METHODS

### Human study population

During the last decade (i.e., January 2003 to May 2021), we have screened 781 individuals for HSV-1 and HSV-2 seropositivity (**Table I**). Five hundred forty-three individuals were White, 238 were non-White (African, Asian, Hispanic and others), 395 were females, and 386 were males. Among these, a cohort of 283 immuno-competent individuals, with an age range of 21-67 (median 30), and who were seropositive for HSV-1 and seronegative for HSV-2, were enrolled in the present study. All patients were negative for HIV, HBV and had no history of immunodeficiency. 680 patients were HSV-1, HSV-2, or HSV-1/HSV-2 seropositive, among which 711 patients were healthy and ASYMP (individuals who, in the absence of therapy, have never had any recurrent herpes disease, ocular, genital or elsewhere, based on their self-report and physician examination. Even a single episode of any herpetic disease in their life span will exclude the individual from this group). The remaining 70 patients were defined as HSV-seropositive symptomatic who suffered with frequent and severe recurrent genital, ocular and/or oro-facial lesions, with two patients having had clinically well-documented repetitive herpes stromal keratitis (HSK) including one patient with ∼20 episodes over 20 yrs. that necessitated several corneal transplantations.

**Table 1:** Cohort of HLA-A*o2:01 positive, HSV seropositive Symptomatic and Asymptomatic Individuals enrolled in this study

| **Subject-level**<br>**Characteristic** | **All Subjects**<br>**(n= 781)** |
| --- | --- |
| **Gender [no.(%)]:** |  |
| Female | 395 (51%) |
| Male | 386 (49%) |
| **Race [no.(%)]:** |  |
| White | 543 (69%) |
| Nonwhite | 238 (31%) |
| Age [median (range) yr.]: | 30 (21-67 yr.) |
| **HSV status [no. (%)]:** |  |
| HSV-1-positive | 283 (36%) |
| HSV-2-positive | 366 (47%) |
| HSV-1- & 2-positive | 31 (4%) |
| HSV-negative | 92 (12%) |
| **HLA [no.(%)]** |  |
| HLA-A*02:01-positive | 411 (53%) |
| HLA-A*02:01-negative | 370 (447%) |
| **Herpes Disease Status [no.(%)]** |  |
| ASYMPTOMATIC (ASYMP) | 711 (91%) |
| SYMPTOMATIC (SYMP) | 70 (9%) |

Signs of recurrent disease in SYMP patients were defined as herpetic lid lesions, herpetic conjunctivitis, dendritic or geographic keratitis, stromal keratitis, and iritis consistent with HSK, with one or more episodes per year for the past 2 years. However, at the time of blood collection, SYMP patients had no recurrent disease (other than corneal scarring) and had no recurrences during the past 30 days. They had no ocular disease other than HSK; had no history of recurrent genital herpes; and are HSV-1 seropositive and HSV-2 seronegative. Because the spectrum of recurrent ocular herpetic disease is wide, our emphasis is mainly on the number of recurrent episodes and not on the severity of the recurrent disease. No attempt was made to assign specific T cell epitopes to specific severity of recurrent lesions. Patients are also excluded if they: (1) have an active ocular (or elsewhere) herpetic lesion or had one in the past 30 days; (2) are seropositive for HSV-2; (3) pregnant or breastfeeding; or (4) were on acyclovir or other related anti-viral drugs or any immunosuppressive drugs at time of blood draw. SYMP and ASYMP groups were matched for age, gender, serological status, and race. Eighty-six healthy control individuals were seronegative for both HSV-1 and HSV-2 and had no history of ocular herpes, genital lesions, or oro-facial herpes disease. All subjects were enrolled at the University of California Irvine under approved Institutional Review Board-approved protocols (IRB# 2003-3111 and IRB#2009-6963). A written informed consent was received from all participants prior to inclusion in the study.

### HLA typing

HLA-A2 sub-typing was performed using a commercial Sequence-Specific Primer (SSP) kit and following the manufacturer’s instructions (SSPR1-A2; One Lambda, Canoga Park, CA). Briefly, genomic DNA extracted from PBMC of HSV-seropositive SYMP and ASYMP individuals was analyzed using a TECAN DNA workstation from a 96-well plate with 2μl volume per well, as we previously described (22). The yield and purity of each DNA sample was tested using a UV-spectrophotometer. The integrity of DNA samples was ascertained by electrophoresis on agarose gel. Each DNA sample was then subjected to multiple small volume PCR reactions using primers specific to areas of the genome surrounding the single point mutations associated with each allele. Only primers that matched the specific sequence of a particular allele would amplify a product. The PCR products were subsequently electrophoresed on a 2.5% agarose gel with ethidium bromide and the pattern of amplicon generation was analyzed using HLA Fusion Software (One Lambda, Inc., Canoga Park, CA). In addition, the HLA-A2 status was confirmed by staining PBMC with anti-HLA-A2 mAb, BB7.2 (BD Pharmingen, USA), at 4°C for 30 min. The cells were washed, acquired on a BD LSR II and analyzed using FlowJo software version 9.7.5 (TreeStar).

### Peripheral blood mononuclear cell (PBMC) isolation

Individuals (negative for HIV, HBV and with or without any HSV infection history) were recruited at Gavin Herbert Eye Institute and UCI Institute for Clinical and Translational Science (ICTS). Between 40 and 100 mL of blood were drawn into yellow-top Vacutainer^®^ Tubes (Becton Dickinson, USA). The serum was isolated and stored at −80°C for detection of anti-HSV-1 and HSV-2 antibodies, as we previously described (8). PBMC were isolated by gradient centrifugation using leukocyte separation medium (Cellgro, USA). The cells were washed in PBS and re-suspended in complete culture medium consisting of RPMI-1640, 10% FBS (Bio-Products, Woodland, CA, USA) supplemented with 1x penicillin/L-glutamine/streptomycin, 1x sodium pyruvate, 1x non-essential amino acids, and 50 μM of 2-mercaptoethanol (Life Technologies, Rockville, MD, USA). Freshly isolated PBMC were also cryo-preserved in 90% FCS and 10% DMSO in liquid nitrogen for future testing.

### Flow cytometry analysis

PBMC were analyzed by flow cytometry after staining with fluorochrome-conjugated human and mouse specific monoclonal antibodies (mAbs). The following anti-human antibodies were used: CD3 (clone SK7) PE-Cy7, CD44 (clone G44-26) A700, CD8 (clone SK1) APC-Cy7, T-bet (clone O4-46) Alexa Fluor 488 (BD Pharmingen); Ki-67 (clone 20Raj1) PE-Cy7, Eomes (clone WD1928) eFluor 660 (eBioscience). CD28 (clone CD28.2) A700; CD57 (clone HCD57) FITC; KLRGI (clone 2F1/KLRG1) APC (BioLegend). gB tetramers were made by NIH Tetramer Core Facility, Emory University, Atlanta, GA. For surface stain mAbs against various cell markers were added to a total of 1×10^6^ PBMC in phosphate-buffered saline containing 1% FBS and 0.1% sodium azide (FACS buffer) for 45 min at 4°C. After washing with FACS buffer, cells were permeabilized for 20 min on ice using the Cytofix/Cytoperm Kit (BD Biosciences) and then washed twice with Perm/Wash Buffer (BD Bioscience). Intra-cellular cytokine/intra-nuclear transcription factor-staining mAbs were then added to the cells and incubated for 45 min on ice in the dark. Cells were washed again with Perm/Wash and FACS Buffer and fixed in PBS containing 2% paraformaldehyde (Sigma-Aldrich, St. Louis, MO). For each sample, 200,000 total events were acquired on BD LSRII. Antibody capture beads (BD Biosciences) were used as individual compensation tubes for each fluorophore in the experiment. To define positive and negative populations, we employed fluorescence minus controls for each fluorophore used in this study, when initially developing staining protocols. In addition, we further optimized gating by examining known negative cell populations for background level expression. The gating strategy was similar to that used in our previous work (5). Briefly, we gated on single cells, dump^−^ cells, viable cells (Aqua Blue), lymphocytes, CD3^+^ cells, and CD8^+^ cells before finally gating human epitope-specific CD8^+^ T cells using gB epitope-specific tetramers. Data analysis was performed using FlowJo version 9.7.5 (TreeStar, Ashland, OR). Statistical analyses were done using GraphPad Prism version 5 (La Jolla, CA).

### CD107 cytotoxicity assay

To detect gB-specific cytolytic CD8^+^ T cells in PBMC, intracellular CD107^a/b^ cytotoxicity assay was performed as described by Betts *et. al*. (23) with a few modifications. Briefly, 1×10^6^ PBMCs from patients were transferred into 96-well flat bottom plates and stimulated with immunodominant and subdominant gB peptide epitopes (10μg/ml in 200μl complete culture medium) in the presence of anti-CD107^a^-FITC, CD107^b^-FITC, and BD Golgi stop (10μg/ml) for 6 hours at 37°C. PHA (10μg/mL) (Sigma) and no peptide were used as positive and negative controls, respectively. At the end of the incubation period the cells were transferred to 96 well round bottom plate and washed once with FACS buffer. Anti-CD107^a^-FITC and anti-CD107^b^-FITC were used for surface stain and IFN-γ was detected using intra-cellular staining as described above.

### Statistical analyses

Data for each assay were compared by analysis of variance (ANOVA) and Student’s *t* test using Graph Pad Prism 5 software (San Diego, CA). Differences between the groups were identified by ANOVA, and multiple comparison procedures, as we previously described (24). Data are expressed as the mean + SD. Results were considered statistically significant at *p* < 0.05.

## RESULTS

### 1. Decline of HSV-specific CD8^+^ T cells in seropositive SYMP individual

The characteristics of the SYMP and ASYMP study population used in the present study, with respect to gender, age, HLA-A*02:01 frequency distribution, HSV-1/HSV-2 seropositivity and status of ocular herpetic disease are presented in **Table I** and detailed in the *Materials and Methods*. Since HSV-1 is the main cause of ocular herpes, only individuals who are HSV-1 seropositive and HSV-2 seronegative were enrolled in the present study. The HSV-1 seropositive individuals were segregated into two groups: (*i*) HLA-A*02:01 positive, HSV-1-infected ASYMP individuals who have never had any clinically detectable herpes disease; and (*ii*) HLA-A*02:01 positive HSV-1-infected SYMP individuals with a history of numerous episodes of well-documented recurrent clinical herpes diseases, such as herpetic lid lesions, herpetic conjunctivitis, dendritic or geographic keratitis, stromal keratitis, and iritis consistent with rHSK, with 1 or more episodes per year for the past 2 years. Only SYMP patients who were not on acyclovir or other anti-viral or anti-inflammatory drug treatments at the time of blood sample collections have been enrolled.

We followed the frequency of CD8^+^ T cells specific to gB_183-191_ epitope, using HLA-A*02:01 specific tetramers/anti-CD8 mAbs, for more than 3 years in the peripheral blood of two HLA-A*02:01 positive, HSV-1 seropositive ASYMP and two HLA-A*02:01 positive, HSV-1 seropositive SYMP individuals (**Fig. 1**). At each time point, history of recurrent disease was recorded. Arrow shows the frequency of recurrent disease. Frequency of gB_183-191_-specific CD8^+^ T cells are shown as FACS dot plots **(Fig. 1A)**. Top panel show dot plot of gB_183-191_-specific CD8^+^ T cells at different time points of representative ASYMP individual. Bottom panel show dot plots of gB_183-191_-specific CD8^+^ T cells at different time points of representative SYMP individual. Line graph **(Fig 1B)** shows the kinetic of gB_183-191_-specific CD8^+^ T cells at different time points of 2 ASYMP individuals. In both ASYMP individuals, frequency of the gB_183-191_-specific CD8^+^ T cells was maintained over a period of time. In one ASYMP individual, frequency of the gB_183-191_-specific CD8^+^ T cells was very similar (4.9% to 5.9%) over a period of time. However, in one ASYMP individual, initially there was a drop in the frequency of the gB_183-191_-specific CD8^+^ T cells (from 5.1% to 3.2%), which was subsequently maintained for another two years (3.2% to 3.8%). Line graph **(Fig. 1C)** shows the kinetic of gB_183-191_-specific CD8^+^ T cells at different time point of two representative SYMP individuals. Unlike ASYMP individuals, frequency of the gB_183-191_-specific CD8^+^ T cells was declined over a period of time. In one SYMP individual, there was a sharp decline in gB_183-191_-specific CD8^+^ T cells (5.3% to 1.8%). However, in another SYMP individual, there was a moderate decline in the gB_183-191_-specific CD8^+^ T cells (from 5.4% to 4.2%). Each arrow on the line graph shows episode of the recurrent disease in SYMP individuals. We next determined the frequency of HSV-specific senescent CD8^+^ T cells by measuring the expression of CD57 PMBC isolated from HSV-1 seropositive ASYMP and SYMP individuals. We also found that the gB_183-191_-specific CD8^+^ T cells from SYMP individuals were exhausted. **Fig. 1D** show representative histogram of PD-1 expression on gB_183-191_-specific CD8^+^ T cells (filled histogram represent expression of PD-1 in SYMP individuals and open histogram represent ASYMP individual). Expression level (**Fig 1E**) and absolute number of (**Fig 1F**) of PD-1 was significantly high in SYMP individuals as compared to ASYMP individuals.

**Figure 1.**
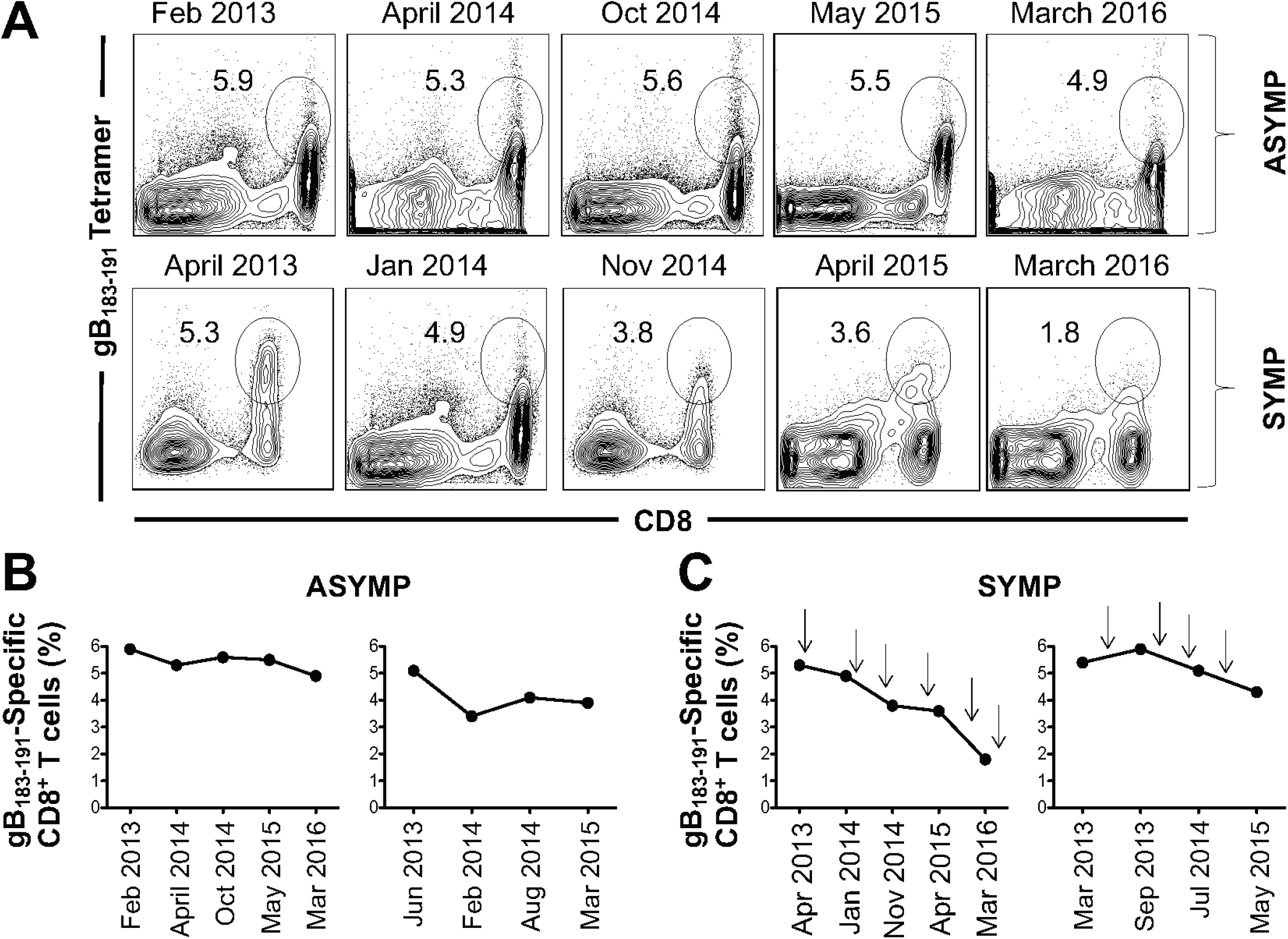
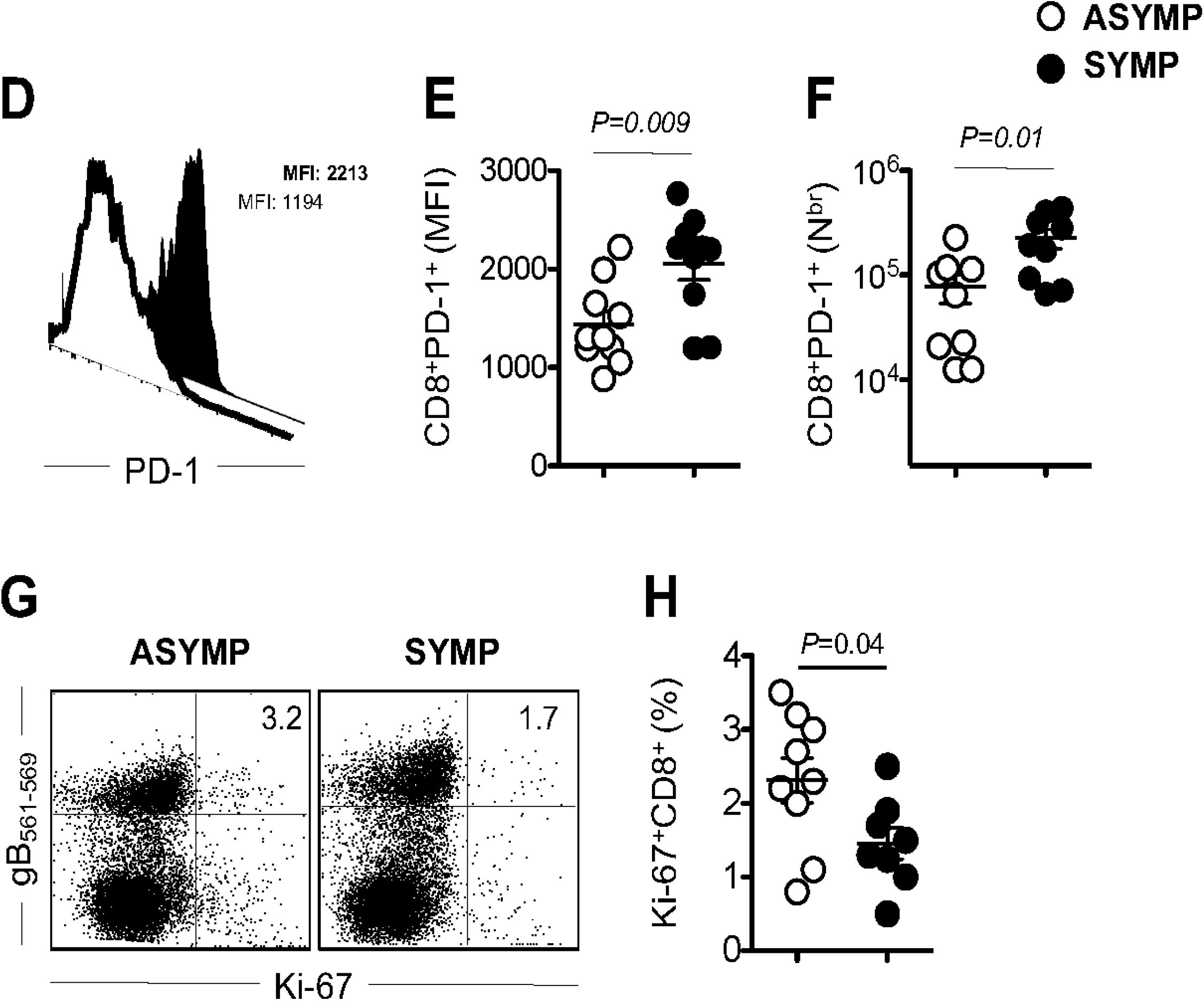
Decline of gB_183-191_-specific CD8^+^ T cells in SYMP individual but not in ASYMP individuals. gB_183-191_-specific CD8^+^ T cells were measured in HLA-A*02:01-positive, HSV-1 seropositive SYMP and ASYMP individuals at different time points. Each time history of recurrent disease was recorded. **(A)** Frequency of gB_183-191_-specific CD8^+^ T cells are shown as FACS dot plots. Top panel show dot plot of gB_183-191_-specific CD8^+^ T cells at different time point of ASYMP individual. Bottom panel show dot plot of gB_183-191_-specific CD8^+^ T cells at different time point of SYMP individual. **(B)** Line graph shows the kinetic of gB_183-191_-specific CD8^+^ T cells at different time point of ASYMP individual. Frequency of the gB_183-191_-specific CD8^+^ T cells was maintained over a period of time. **(C)** Line graph shows the kinetic of gB_183-191_-specific CD8^+^ T cells at different time point of SYMP individual. Unlike ASYMP individuals, frequency of the gB_183-191_-specific CD8^+^ T cells was declined over a period of time. Arrow on the line graph shows the frequency of recurrent disease in SYMP individuals.

Further we evaluated the proliferative capacity of gB_183-191_-specific CD8^+^ T cells from SYMP and ASYMP individuals using intra-nuclear staining of Ki-67 to see if there is any difference in the recent proliferation of CD8^+^ T cells. Representative dot plot of gB_183-191_-specific CD8^+^ T cells stained for proliferation marker Ki-67 (**Fig. 1G**). We looked at the frequency of Ki-67^+^CD8^+^ T cells gated on gB_183-191_ tetramer and found that there was significantly (*P = 0*.*04*) less proliferation of CD8^+^ T cells isolated from SYMP vs. ASYMP individuals. In SYMP individual median frequency of Ki-67^+^CD8^+^ T was recorded as 2.3% whereas in ASYMP individual it was recorded as 1.3% (**Fig. 1H**). Open circles indicate data of ASYMP individuals and filled circle represent data of SYMP individuals. The results are representative of two independent experiments for each individual. Altogether these data clearly indicate that there is higher frequency of terminally differentiated gB_183-191_-specific CD8^+^ T cells which has lost proliferative capacity in SYMP than ASYMP individuals. Unlike ASYMP individuals, HSV-specific CD8^+^ T cells from SYMP individual were not maintained and functionally exhausted. Next, we wanted to see if there is any possibility of senescence, which might cause decline in the frequency of CD8^+^ T cells in SYMP individuals but not in ASYM individuals.

### 2. Increased frequency of senescent CD8^+^ T cells in HSV-seropositive SYMP individual compared to ASYMP individuals

Next, we investigated if there is any role of senescent CD8^+^ T cell in SYMP vs. ASYMP individuals. PBMC isolated from HLA-A*02:01-positive, HSV-1 seropositive SYMP and ASYMP individuals were stained for senescent marker CD57 on HSV specific CD8^+^ T cells. **Fig 2A** is representative histogram of CD57 expression on gB_183-191_-specific CD8^+^ T cells from ASYMP and SYMP individuals. Right panel show the overlap of CD57 expression from ASYMP and SYMP individuals. Frequency of CD8^+^CD57^+^ on gB_183-191_-specific CD8^+^ T cells from ASYMP and SYMP individuals are shown in **Fig 2B**. There is significantly (*P=0*.*001*) higher percentage of gB_183-191_-specific CD8^+^ T cells expressing CD57 in SYMP individuals than ASYMP individuals. Median expression of CD57 on gB_183-191_-specific CD8^+^ T cells isolated from SYMP individual was 42.3% whereas it was 18.1% in ASYMP individuals. Absolute number of CD8^+^CD57^+^ on gB_183-191_-specific CD8^+^ T cells are shown in **Fig 2C**. We observed that there was significantly (*P=0*.*002*) higher number of gB_183-191_-specific CD8^+^ T isolated from SYMP than ASYMP individuals. As shown in **Fig 2D**, we did not observe significant difference in the Mean Fluorescent Intensity (MFI) of CD57 expression gated on gB_183-191_-specific CD8^+^ T cells from SYMP and ASYMP individuals. However increased level of CD57 expression was observed in SYMP individuals than ASYMP individuals. Open circles indicate data of ASYMP individuals and filled circle represent data of SYMP individuals. The results are representative of two independent experiments for each individual.

**Figure 2.**
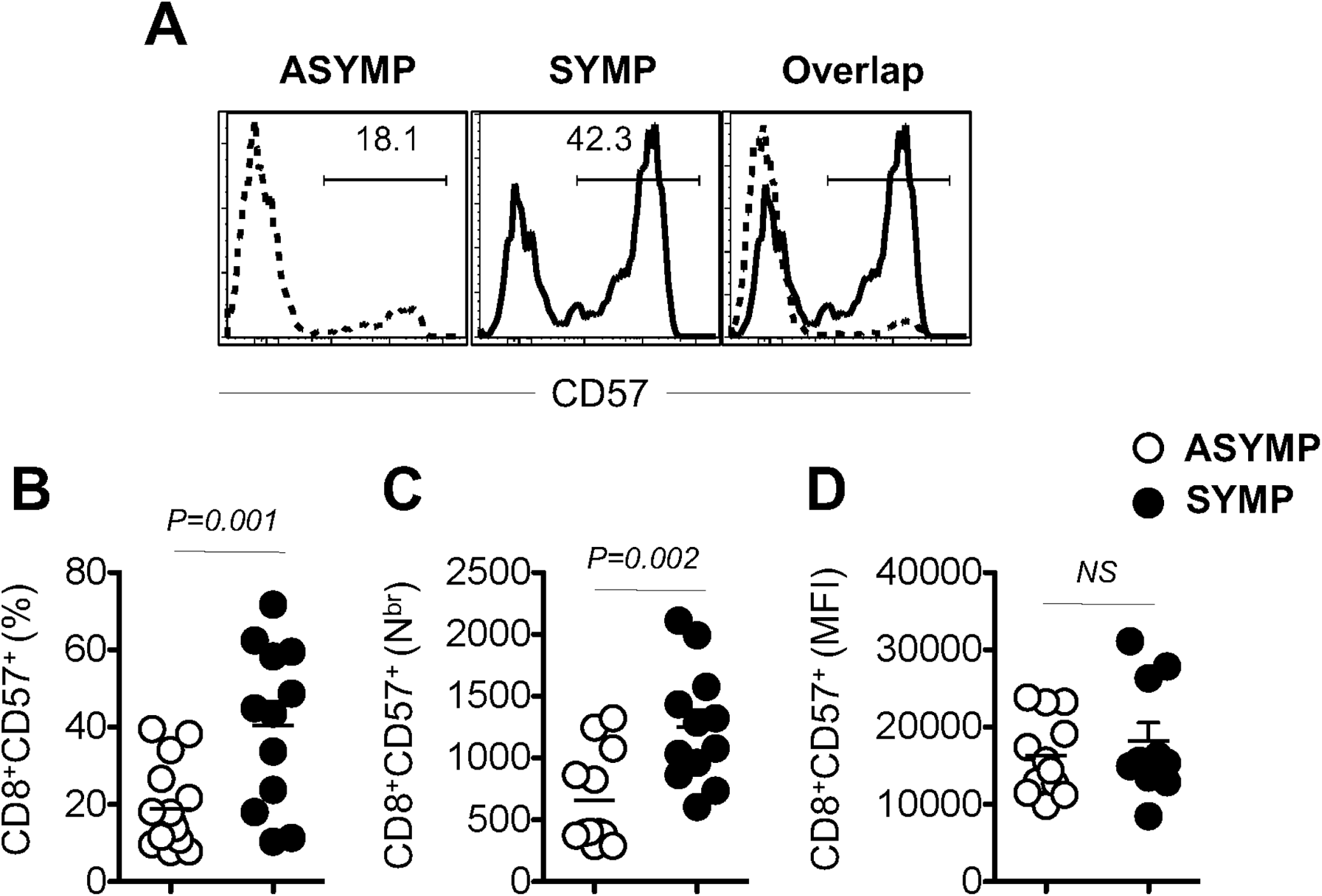
Increased frequency of senescent CD8^+^ T cells in SYMP individual than SYMP individuals. PBMC isolated from HLA-A*02:01-positive, HSV-1 seropositive SYMP and ASYMP individuals were stained for senescent marker CD57. **(A)** Representative histogram of CD57 expression on gB_183-191_-specific CD8^+^ T cells. **(B)** Frequency of CD8^+^CD57^+^ on gB_183-191_-specific CD8^+^ T cells. **(C)** Absolute number of CD8^+^CD57^+^ on gB_183-191_-specific CD8^+^ T cells. **(D)** Mean Fluorescent Intensity (MFI) of CD57 expression gated on gB_183-191_-specific CD8^+^ T cells. Open circles indicate data of ASYMP individuals and filled circle represent data of SYMP individuals. The results are representative of two independent experiments for each individual. The indicated *P* values, calculated using one-way ANOVA, show statistical significance of differences between SYMP and ASYMP individuals. The data are representative of two independent experiments and the bars represent SD between the experiments.

Next, we wanted to visualize the expression of CD57 on gB_183-191_-specific CD8^+^ T from SYMP and ASYMP individuals using powerful tool of capturing images stained with fluorescent mAbs (ImageStream). PBMC were stained with different surface markers (KLRG-1, gB_183-191_ tetramer, CD57 and CD8) similar to FACS staining as outlined in materials and methods section. Image of each cell was captured using ImageStream (EMD Millipore) and data was analyzed by software IDEA V4. **Fig. 3** shows the image of individual cell showing expression of KLRG-1, gB_183-191_ tetramer, CD57, CD8 and superimposed image of all 4 mAbs on PBMC isolated from HSV-1 seropositive ASYMP (**Fig. 3A**) and SYMP individual (**Fig. 3B**). As shown in ImageStream, the expression of CD57 was higher in PBMC isolated from SYMP than ASYMP individuals. However, there were no differences in the staining of gB_183-191_ tetramer and CD8^+^ T cells in PBMC isolated from both groups. Altogether we found that the expression of CD57 was significantly high in SYMP individuals as compared to ASYMP individuals.

**Figure 3.**
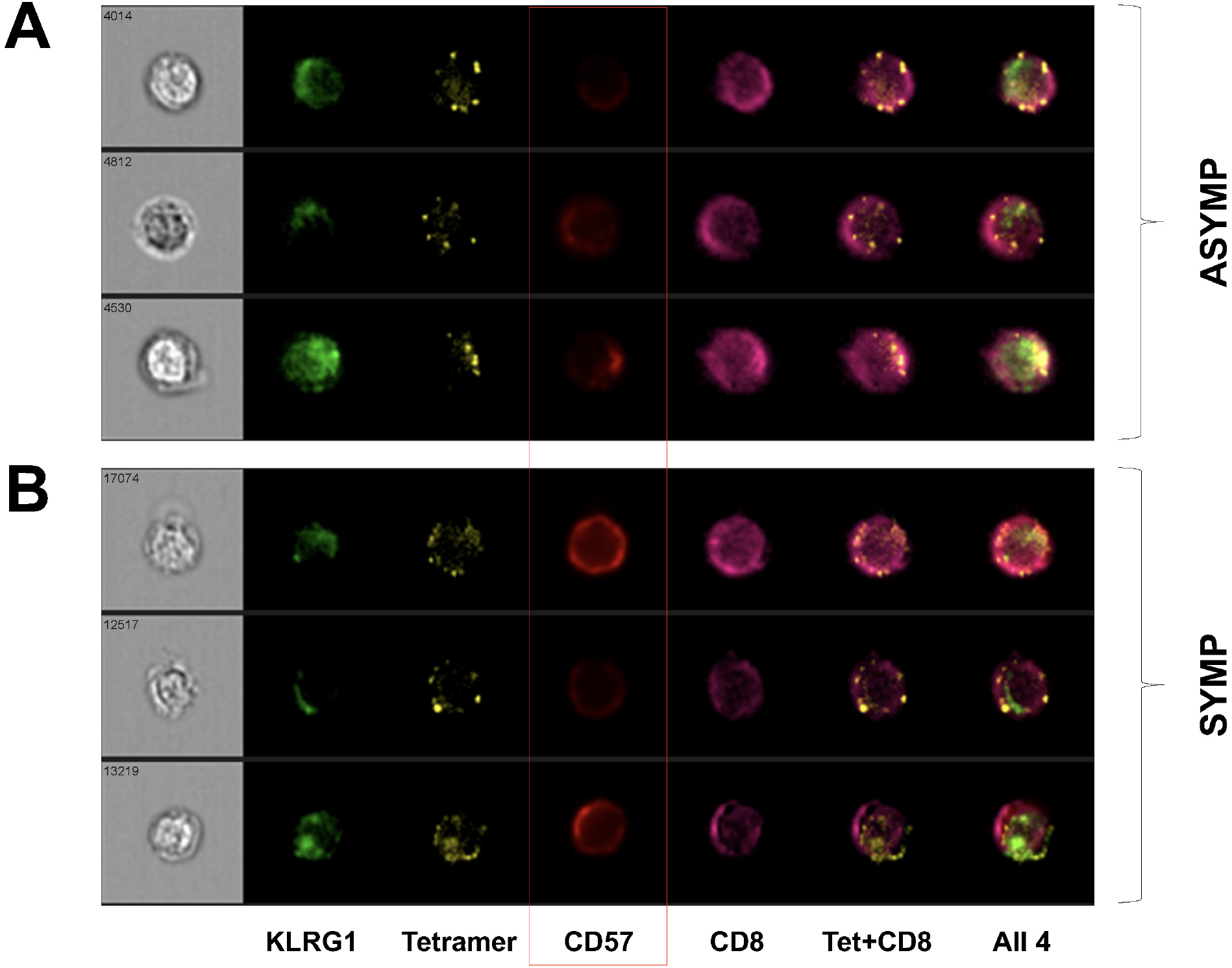
Increased expression of CD57 on gB_183-191_-specific CD8^+^ T cells in SYMP individual than SYMP individuals. PBMC isolated from HLA-A*02:01-positive, HSV-1 seropositive SYMP and ASYMP individuals were stained for senescent marker CD57 along with, CD8, gB_183-191_ tetramer and KLRG-1. Images of individual PBMC stained with different markers were visualized and captured by ImageStream. Data was analyzed using software IDEA version 4. **(A)** Representative images of individual PMBC isolated from HSV-1 seropositive SYMP patient. Cells are shown as individually stained with KLRG-1, gB_183-191_ tetramer, CD57, CD8, gB_183-191_ tetramer and CD8, and superimposed image of PBMC stained with KLRG-1, gB_183-191_ tetramer, CD57 and CD8^+^ T cells.

### 3. HSV-specific CD8^+^ T cells isolated from SYMP individuals are terminally differentiated

We next studied the expression of markers expressed on terminally differentiated gB_183-191_ tetramer specific CD8^+^ T cells. Since terminally differentiated CD8^+^ T cells do not proliferate so we stained cells with Ki-67 to study proliferative capacity. Representative contour plot of KLRG-1^+^CD57^+^ expression on terminally differentiated gB_183-191_-specific CD8^+^ T cells isolated from SYMP and ASYMP individuals are shown in **Fig. 4A**. Frequency of KLRG-1^+^CD57^+^ on gB_183-191_-specific CD8^+^ T cells isolated from SYMP individual (37.9%) was significantly (*P = 0*.*006*) high as compared to ASYMP individual (12.4%) as shown in **Fig. 4B**. Next, we looked at the absolute number of gB_183-191_-specific CD8^+^ T cells expressing KLRG-1^+^CD57^+^ isolated from SYMP and ASYMP individuals. Absolute number of KLRG-1^+^CD57^+^ expressing gB_183-191_-specific CD8^+^ T cells was high on SYMP as compared to ASYMP individual (**Fig. 4C)**. Though the difference in absolute number was not significantly different in SYMP vs. ASYMP individuals. PBMC isolated from HLA-A*02:01-positive, HSV-1 seropositive SYMP and ASYMP individuals were stained for transcription factor T-bet along with the marker of senescent CD57 on gB_183-191_-specific CD8^+^ T cells isolated from SYMP and ASYMP individuals. Representative contour plot of T-betHi^+^CD57Hi^+^ gated on gB_183-191_-specific CD8^+^ T cells in PBMC isolated from SYMP and ASYMP individuals (**Fig. 4D**). We observed significant difference (*P = 0*.*02*) in the frequency of T-bet^Hi^CD57^Hi^ gated on gB_183-191_-specific CD8^+^ T cells isolated from SYMP as compared to ASYMP individuals. Median frequency of T-bet^Hi^CD57^Hi^ gated on gB_183-191_-specific CD8^+^ T cells in SYMP individual was recorded as 26.6% whereas in ASYMP individual it was recorded as 10.2% (**Fig. 4E**). We also observed significant (*P = 0*.*03*) difference in the absolute number of T-bet^Hi^CD57^Hi^ gated on gB_183-191_-specific CD8^+^ T cells. SYMP individual harbor higher absolute number of T-bet^Hi^CD57^Hi^ gated on gB_183-191_-specific CD8^+^ T cells as compared to ASYMP individuals (**Fig. 4F**). Open circles indicate data of ASYMP individuals and filled circle represent data of SYMP individuals. The results are representative of two independent experiments for each individual.

**Figure 4.**
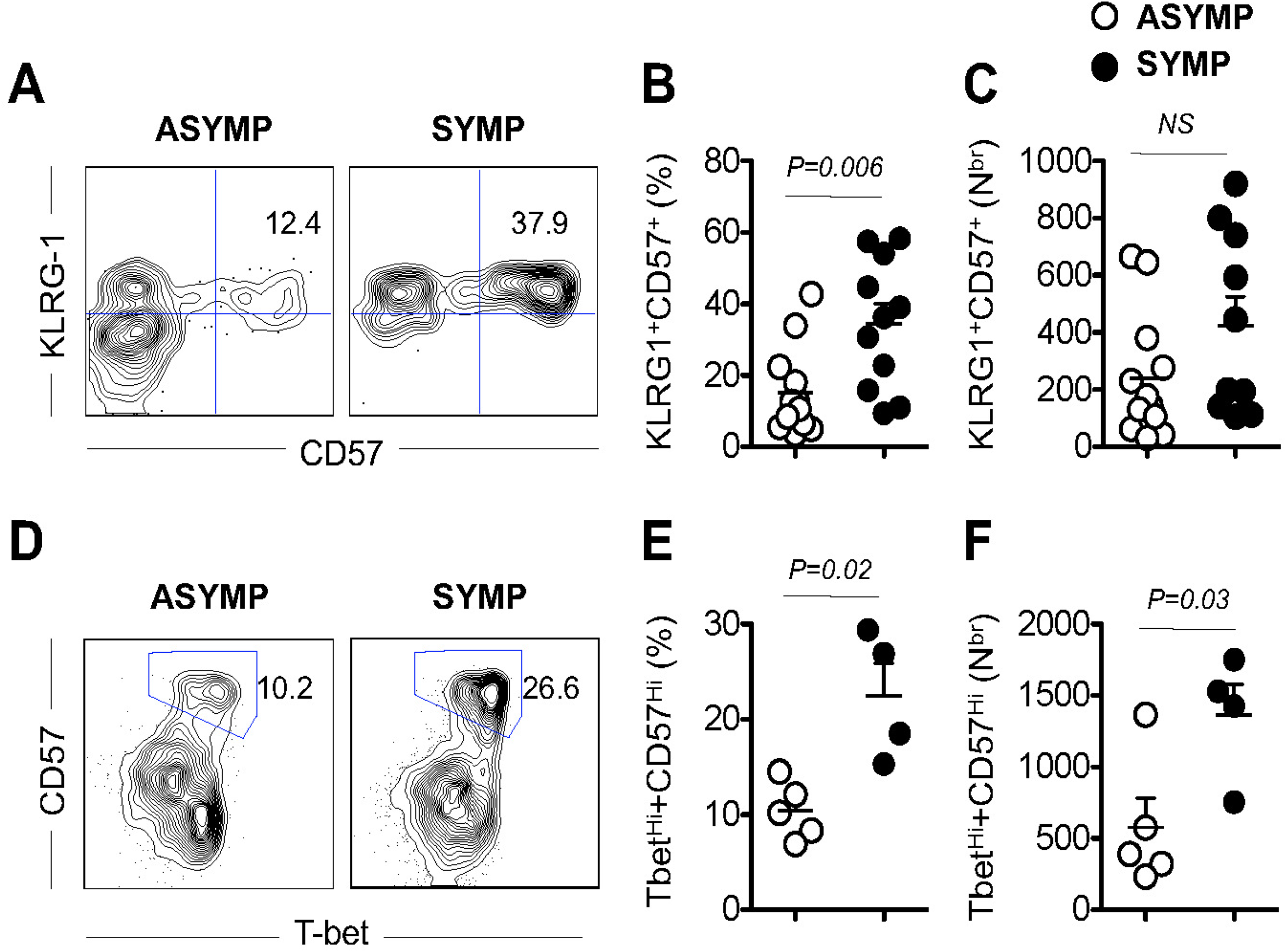
HSV-1 specific CD8^+^ T cells from SYMP individuals are terminally differentiated and lack ability to divide. PBMC isolated from HLA-A*02:01-positive, HSV-1 seropositive SYMP and ASYMP individuals were stained for terminally differentiated marker (KLRG-1) and CD57. Cells were also stained for Ki-67 to see the percentage of proliferating cells. **(A)** Representative contour plot of CD57 expression on terminally differentiated CD8^+^ T cells gated on gB_183-191_ tetramer in PBMC isolated from SYMP and ASYMP individuals. **(B)** Frequency of KLRG-1^+^CD57^+^ on gB_183-191_-specific CD8^+^ T cells. **(C)** Absolute number of KLRG-1^+^CD57^+^ on gB_183-191_-specific CD8^+^ T cells. **(D)** Representative dot plot of gB_183-191_-specific CD8^+^ T cells stained for proliferation marker Ki-67. **(E)** Frequency of Ki-67^+^CD8^+^ T cells gated on gB_183-191_ tetramer. Open circles indicate data of ASYMP individuals and filled circle represent data of SYMP individuals. The results are representative of two independent experiments for each individual. The indicated *P* values, calculated using one-way ANOVA, show statistical significance of differences between SYMP and ASYMP individuals. The data are representative of two independent experiments and the bars represent SD between the experiments.

### 4. Decreased expression of co-stimulatory molecule CD28 and survival molecule CD127 on PBMC isolated from HSV-seropositive SYMP individuals

PBMC isolated from HLA-A*02:01-positive, HSV-1 seropositive SYMP and ASYMP individuals were stained for co-stimulatory and survival molecules to further evaluate if there is any defect in the homeostatic maintenance of gB_183-191_-specific CD8^+^ T cells in SYMP vs. ASYMP individuals. We studies surface expression of co-stimulatory molecule CD28 and survival molecule CD127 on PBMC isolated from HSV-seropositive SYMP and ASYMP individuals **(Fig. 5)**. Representative contour plot of CD28 and CD57 expression gated on gB_183-191_ tetramer in PBMC isolated from SYMP and ASYMP individuals **(Fig. 5A)**. Frequency of the CD28^+^gB_183-191_-specific CD8^+^ T cells was significantly (*P = 0*.*003*) low in SYMP than ASYMP individuals. Median frequency of CD28^+^gB_183-191_-specific CD8^+^ T cells in SYMP individual was recorded as 54.1% whereas in ASYMP individual it was recorded as 25.6% **(Fig. 5B)**. Further we counted the CD28^+^gB_183-191_-specific CD8^+^ T cells and found that there was significant (*P = 0*.*02%*) difference in the absolute number between SYMP and ASYMP individuals. This clearly indicates that there is low expression of co-stimulatory molecules, which further confirm our finding of low proliferation in CD8^+^ T cells isolated from SYMP individuals **(Fig. 5C)**. The next question, which we wanted to address, was if there is any defect in the homeostatic maintenance of gB_183-191_-specific CD8^+^ T cells in SYMP vs. ASYMP individuals. As expected, we found that there was low expression of survival molecule CD127 on gB_183-191_-specific CD8^+^ T cells isolated from SYMP individual than ASYMP individual. **Fig. 5D** is representative histograms showing expression level of survival molecule CD127 gated on gB_183-191_-specific CD8^+^ T cells. The frequency of CD127 gated on gB_183-191_-specific CD8^+^ T cells was significantly (*P = 0*.*02*) low in SYMP than ASYMP individuals (**Fig. 5E**). We recorded median frequency of CD127^+^gB_183-191_-specific CD8^+^ T cells 90.1% in ASYMP individuals whereas in SYMP individuals, it was 74.9%. Similarly, there was significant (*P = 0*.*04*) difference in the absolute number of CD127^+^gB_183-191_-specific CD8^+^ T cells. SYMP individuals have low absolute number of gB_183-191_-specific CD8^+^ T cells expressing survival molecule as compared to ASYMP individuals (**Fig. 5F**). We did not see significant difference in the Mean Fluorescent Intensity (MFI) of CD127 expression on gB_183-191_-specific CD8^+^ T cells between SYMP and ASYMP individuals (**Fig. 5G**). Open circles indicate data of ASYMP individuals and filled circle represent data of SYMP individuals. The results are representative of two independent experiments for each individual.

**Figure 5.**
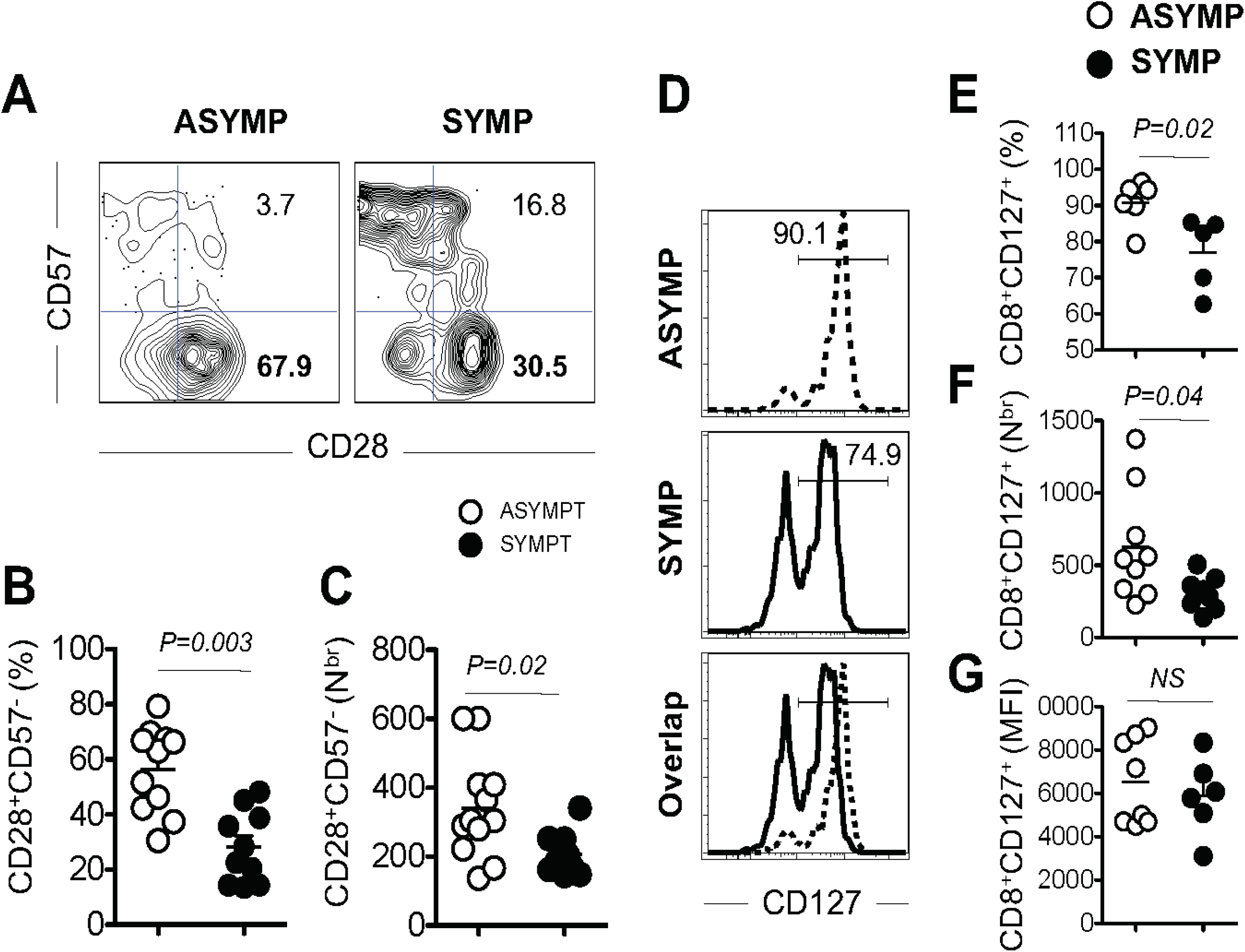
Decreased expression of co-stimulatory molecule CD28 and survival molecule CD127 on PBMC isolated from SYMP individuals. PBMC isolated from HLA-A*02:01-positive, HSV-1 seropositive SYMP and ASYMP individuals were stained for co-stimulatory and survival molecules. **(A)** Representative contour plot of CD28 and CD57 expression gated on gB_183-191_ tetramer in PBMC isolated from SYMP and ASYMP individuals. **(B)** Frequency of CD28^+^CD57^-^ on gB_183-191_-specific CD8^+^ T cells. **(C)** Absolute number of CD28^+^CD57^-^ on gB_183-191_-specific CD8^+^ T cells. **(D)** Representative histogram showing expression level of survival molecule CD127 gated on gB_183-191_-specific CD8^+^ T cells. **(E)**. Frequency of CD127 gated on gB_183-191_-specific CD8^+^ T cells. **(F)** Absolute number of CD127 gated on gB_183-191_-specific CD8^+^ T cells. **(G)** Mean Fluorescent Intensity (MFI) of CD127 expression on gB_183-191_-specific CD8^+^ T cells. Open circles indicate data of ASYMP individuals and filled circle represent data of SYMP individuals. The results are representative of two independent experiments for each individual. The indicated *P* values, calculated using one-way ANOVA, show statistical significance of differences between SYMP and ASYMP individuals. The data are representative of two independent experiments and the bars represent SD between the experiments.

### 5. Decreased production of effector molecules Granzyme B and Perforin by HSV-specific CD8+ T cells isolated from HSV-1 seropositive SYMP individuals

PBMC isolated from HLA-A*02:01-positive, HSV-1 seropositive SYMP and ASYMP individuals were stained for effector molecules granzyme B and perforin after *in vitro* stimulation. (**Fig 6**). Representative contour plot of CD8^+^GzmB^+^ cells gated on gB_183-191_-specific CD8^+^ T cells in PBMC isolated from SYMP and ASYMP individuals. We observed significant difference (*P = 0*.*006*) in the frequency of CD8^+^GzmB^+^ gated on gB_183-191_-specific CD8^+^ T cells isolated from SYMP as compared to ASYMP individuals (**Fig 6B**). Median frequency of CD8^+^GzmB^+^ gated on gB_183-191_-specific CD8^+^ T cells in SYMP individual was recorded as 17.4% whereas in ASYMP individual it was recorded as 29.1%. We also observed significant (*P = 0*.*03*) difference in the absolute number of CD8^+^GzmB^+^ gated on gB_183-191_-specific CD8^+^ T cells. ASYMP individual harbor higher absolute number of CD8^+^GzmB^+^ gated on gB_183-191_-specific CD8^+^ T cells as compared to SYMP individuals (**Fig 6C**). We also looked at the production of another effector molecule Perforin on HSV-specific CD8^+^ T cells. Representative contour plot of CD8^+^Perforin^+^ cells gated on gB_183-191_-specific CD8^+^ T cells in PBMC isolated from SYMP and ASYMP individuals (**Fig. 6D**). We observed significant difference (*P = 0*.*03*) in the frequency of CD8^+^Perforin^+^ gated on gB_183-191_-specific CD8^+^ T cells isolated from SYMP as compared to ASYMP individuals (**Fig 6E**). Median frequency of CD8^+^Perforin^+^ gated on gB_183-191_-specific CD8^+^ T cells in SYMP individual was recorded as 10.7% whereas in ASYMP individual it was recorded as 19.8%. We also observed significant (*P = 0*.*02*) difference in the absolute number of CD8^+^Perforin^+^ gated on gB_183-191_-specific CD8^+^ T cells. ASYMP individual harbor higher absolute number of CD8^+^Perforin^+^ gated on gB_183-191_-specific CD8^+^ T cells as compared to SYMP individuals (**Fig 6F**). Open circles indicate data of ASYMP individuals and filled circle represent data of SYMP individuals. The results are representative of two independent experiments for each individual. Altogether, these results indicate that, there is significantly higher frequency of HSV specific senescent CD8^+^ T cell in SYMP individuals, which are exhausted, lack proliferative capacity, decreased expression of co-stimulatory molecule, less functional and cannot maintain homeostatic proliferation as compared to HSV-seropositive ASYMP individuals.

## DISCUSSION

Immunosenescence is marked by a progressive increase in the number of memory CD8^+^ T cells showing poor functionality in terms of killing persistent virus. In this report we put forward a possibility of immunosenescence, which might lead to poor functionality of CD8^+^ T cell cytotoxicity against reactivating virus in SYMP individuals whereas in ASYMP individuals CD8^+^ T cell cytotoxicity is intact. The other question, which we wanted to address, is there any difference in the maintenance of optimal CD8^+^ T cell response between HSV-1 seropositive ASYMP and SYMP individuals. Senescent CD8^+^ T cells appear to preserve a limited degree of responsiveness to antigenic stimulation in spite of defective replication abilities. These CD8^+^ T cells express higher levels of markers linked to T cell activation (25), senescence and immune exhaustion (26) and are deficient in markers involved in co-stimulation (27) and T-cell survival. Immune activation in HIV-infected individuals significantly decreased expression of CD127 and increased CD57, which brings evidence of a T cell subset with limited renewing capacity and survival competence (28). CD57 is a marker of senescence, expressed by most terminally differentiated CD8^+^ T cells. The rate of turnover of CD57^+^ CD8^+^ T cells has been shown to increase in conditions associated with immune dysregulation and autoimmune diseases (16). In concordance with other studies involving HIV-infected T cells expressing CD57 (29), here we showed that HSV-specific CD8^+^ T cells express increased levels of CD57 along with decreased expression of CD28, suggesting the onset of senescence stimulated *ex vivo* by reactivating virus. CD57 and KLRG1 are increased with age and correlate with effector memory subsets, however several groups have associated CD57 and KLRG1 with loss of function (16, 30, 31). A decreased proportion of CD28^+^ CD8^+^ T cells in SYMP individuals represent another evidence for immunosenescence. Chronic HCV infection causes a contraction in early differentiated CD28^+^ HCV and HIV-specific T cells, which indicate that virus-infected cytotoxic T cells reached a state of replicative senescence (13, 32, 33). Our data also suggest that HSV infection in SYMP individuals enhances the differentiation of CD8^+^ T cells towards a late-differentiation phenotype, which could be defective in virus elimination. We found that the expression of CD127 was low on HSV-specific CD8^+^ T cells isolated from SYMP individuals as compared to ASYMP individuals. Others have shown that the progressive loss of CD127 in chronic HIV infections leads to increased apoptosis of CD8^+^ T cells (34). Our data showed that there was significantly higher frequency of HSV-specific CD57^+^CD8^+^ T cells in SYMP individuals as compared to ASYMP individuals. This data is in accordance with increased expression of CD57 was associated with high level of KLRG1 and low level of CD28 is characteristic of senescent CD8^+^ T cells (20). The T-box transcription factor T-bet plays crucial roles in determining differential fate of CD8^+^ T cells responding to infection, optimal memory, and terminal differentiation. In CD8^+^ T cells, T-bet is up regulated upon activation and is associated with induction of effector functions, including cytotoxicity (35). Expression of T-bet was increased in CD8^+^ T cells from SYMP individuals and correlated closely with expression of CD57 and KLRG1. Although senescent CD8^+^ T cells typically have preserved effector function (i.e., the ability to produce cytokines and even kill target cells), lack of proliferative potential impairs their ability to mount robust immune responses and expand in number upon reactivation (36, 37). In this report we have clearly shown that the intermittent virus reactivation leads to loss of early-differentiated CD8^+^ T cells and progressive accumulation of repeatedly activated, late-differentiated senescent CD8^+^ T cells in SYMP individuals. Our observations on immunological changes occurring on CD8^+^ T cells isolated form SYMP individuals eventually drive them towards the end stage of senescence likely favoring viral persistence.

**Figure 6.**
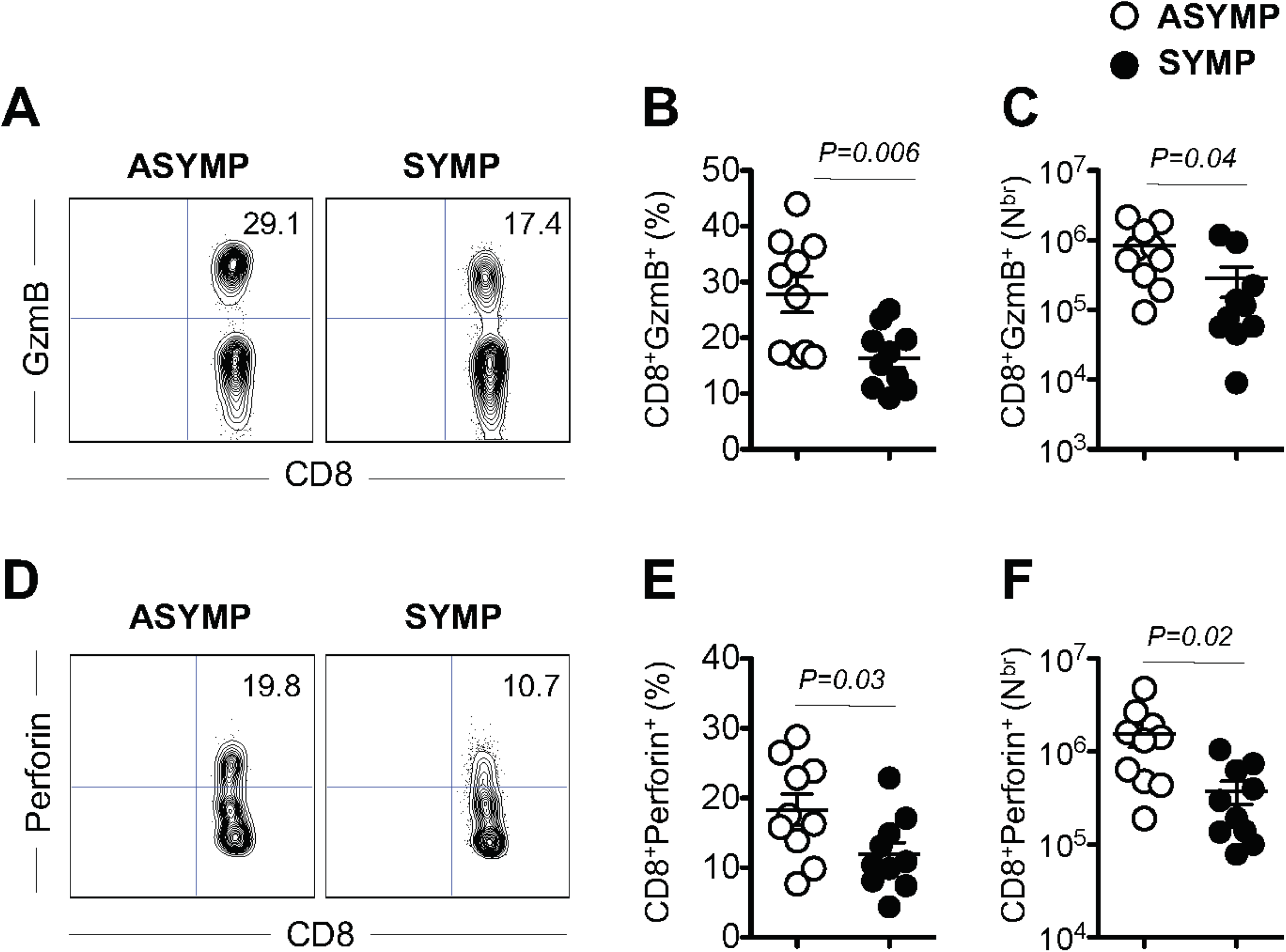
Increased expression of T-bet on terminally differentiated CD8^+^ T cells isolated from SYMP individuals. PBMC isolated from HLA-A*02:01-positive, HSV-1 seropositive SYMP and ASYMP individuals were stained for transcription factor T-bet along with CD57. **(A)** Representative contour plot of T-betHi^+^CD57^+^ expression gated on gB_183-191_ tetramer in PBMC isolated from SYMP and ASYMP individuals. **(B)** Frequency of T-betHi^+^CD57^+^ on gB_183-191_-specific CD8^+^ T cells. **(C)** Absolute number of T-betHi^+^CD57^+^ on gB_183-191_-specific CD8^+^ T cells. Open circles indicate data of ASYMP individuals and filled circle represent data of SYMP individuals. The results are representative of two independent experiments for each individual. The indicated *P* values, calculated using one-way ANOVA, show statistical significance of differences between SYMP and ASYMP individuals. The data are representative of two independent experiments and the bars represent SD between the experiments.

## ACKNOWLEDGEMENTS

This work is supported by Public Health Service Research R01 Grants EY026103, EY019896 and EY024618 from National Eye Institute (NEI) and R21 Grant AI158060, AI150091, AI143348, AI147499, AI143326, AI138764, AI124911 and AI110902 from National Institutes of allergy and Infectious Diseases (NIAID) (to L.BM.), and in part by The Discovery Center for Eye Research (DCER) and the Research to Prevent Blindness (RPB) grant. The authors would like to thank Dale Long from the NIH Tetramer Facility (Emory University, Atlanta, GA) for providing the Tetramers used in this study. We also thank Barbara Bodenhoefer (RN) from UC Irvine’s Institute for Clinical and Translational Science (ICTS) for helping with blood drawing from HSV-1 seropositive symptomatic and asymptomatic individuals.

## Notes

Conflict of interest: The authors have declared that no conflict of interest exists

$ Footnotes: This work is supported by Public Health Service Research R01 Grants EY026103, EY019896 and EY024618 from National Eye Institute (NEI) and R21 Grant AI158060, AI150091, AI143348, AI147499, AI143326, AI138764, AI124911 and AI110902 from National Institutes of allergy and Infectious Diseases (NIAID) (to L.BM.), and in part by The Discovery Center for Eye Research (DCER) and the Research to Prevent Blindness (RPB) grant.

